# Ecology and evolution of chlamydial symbionts of arthropods

**DOI:** 10.1101/2022.03.11.483957

**Authors:** Tamara Halter, Stephan Köstlbacher, Astrid Collingro, Barbara S. Sixt, Elena R. Tönshoff, Frederik Hendrickx, Rok Kostanjšek, Matthias Horn

**Affiliations:** Centre for Microbiology and Environmental Systems Science, University of Vienna; The Laboratory for Molecular Infection Medicine Sweden (MIMS), Umeå Centre for Microbial Research (UCMR), Department of Molecular Biology, Umeå University; Institute of Molecular Biology and Biophysics, Eidgenössische Technische Hochschule Zürich (ETH), Zürich, Switzerland; Royal Belgian Institute of Natural Sciences; Department of Biology, Biotechnical Faculty, University of Ljubljana

## Abstract

The phylum Chlamydiae consists of obligate intracellular bacteria including major human pathogens and diverse environmental representatives. Here we investigated the Rhabdochlamydiaceae, which is predicted to be the largest and most diverse chlamydial family, with the few described members known to infect arthropod hosts. Using published 16S rRNA gene sequence data we identified at least 388 genus-level lineages containing about 14 051 putative species within this family. We show that rhabdochlamydiae are mainly found in freshwater and soil environments, suggesting the existence of diverse, yet unknown hosts. Next, we used a comprehensive genome dataset including metagenome assembled genomes classified as members of the family Rhabdochlamydiaceae, and we added novel complete genome sequences of *Rhabdochlamydia porcellionis* infecting the woodlouse *Porcellio scaber*, and of ‘*Candidatus* R. oedothoracis’ associated with the linyphiid dwarf spider *Oedothorax gibbosus*. Comparative analysis of basic genome features and gene content with reference genomes of well-studied chlamydial families with known host ranges, namely Parachlamydiaceae (protist hosts) and Chlamydiaceae (human and other vertebrate hosts) suggested distinct niches for members of the Rhabdochlamydiaceae. We propose that members of the family represent intermediate stages of adaptation of chlamydiae from protists to vertebrate hosts. Within the genus *Rhabdochlamydia*, pronounced genome size reduction could be observed (1.49-1.93 Mb). The abundance and genomic distribution of transposases suggests transposable element expansion and subsequent gene inactivation as a mechanism of genome streamlining during adaptation to new hosts. This type of genome reduction has never been described before for any member of the phylum Chlamydiae. This study provides new insights into the molecular ecology, genomic diversity, and evolution of representatives of one of the most divergent chlamydial families.

## Introduction

The phylum Chlamydiae was originally regarded as a small group of obligate intracellular bacteria infecting humans and few animal species [1]. Today, the chlamydiae are known to be associated with a broad spectrum of host organisms including protists, arthropods, and diverse vertebrates [2–6]. Some of those may also infect mammalian cells and have thus been proposed to represent emerging human pathogens [7–9]. While cultured representatives of only six families are available to date, molecular surveys suggest that a large undiscovered diversity exists, with over one thousand family-level lineages in various environments worldwide [6, 10].

All chlamydiae share a common ancestor that has lived around one billion years ago, and there is evidence that the emergence of their unique and strictly intracellular lifestyle dates back to these Precambrian times [11–13]. The characteristic biphasic developmental cycle of characterized representatives consists of the infective elementary bodies (EBs) that enter eukaryotic host cells and transform into replicative reticulate bodies (RBs). Inside the host cells, chlamydiae stay in host-derived vacuoles termed inclusions. Eventually, RBs differentiate back to EBs, exit the host cell either by lysis or extrusion and start a new infection cycle [14].

Genomics has helped to gain fundamental insights into chlamydial biology, host adaptation, and evolution. Chlamydiae generally have small, reduced genomes, and lack metabolic pathways that are complemented by importing host cell metabolites [15]. Despite recent advances in genetic manipulation of members of the well-studied family Chlamydiaceae, like *Chlamydia trachomatis* [16, 17], genomics remains of utmost importance to study the more elusive chlamydiae found in the environment, collectively referred to as environmental chlamydiae. For instance, genomics revealed that the chlamydial developmental cycle, including major virulence mechanisms such as the type III secretion system, is well conserved also among the environmental representatives [3, 11, 12, 18]. Yet, the genetic repertoire of environmental chlamydiae is generally more versatile than that of Chlamydiaceae, including more complete metabolic pathways and richer arsenals of predicted effector proteins to interact with their evolutionary distinct eukaryotic host cells [19–21]. More recently, single cell genomics and large-scale metagenomics revealed a surprising biological variability of environmental chlamydiae, including evidence for motility and a widespread potential for anaerobic metabolism [6, 22–24].

The family Rhabdochlamydiaceae is putatively one of the largest and most diverse - yet poorly studied clades - within the phylum Chlamydiae [10]. The only known hosts of rhabdochlamydiae are arthropods including ticks, spiders, cockroaches, and woodlice [25–28]. An infection with rhabdochlamydiae was reported to be detrimental for cockroaches and woodlice, leading to severe abdominal swelling [26, 29] or heavy tissue damage [30] in the respective host. However, the prevalence was reported to be low, accounting to 1 % on average for ticks [31], and to 15 % on average for woodlice [30]. Although rhabdochlamydiae are potentially important members of the phylum, so far there is only one described draft genome sequence available from ‘*Candidatus* Rhabdochlamydia helvetica’ [28]. Here we add the complete genome sequences of *Rhabdochlamydia porcellionis* [25] and the new species ‘*Candidatus* Rhabdochlamydia oedothoracis’ [27], and we use a collection of metagenome assembled genomes (MAGs) to investigate the biology and evolution of members of the Rhabdochlamydiaceae. We provide evidence for a large, yet undiscovered diversity of rhabdochlamydiae especially in freshwater and soil ecosystems. We show that their genomic setup suggests a host spectrum beyond arthropods and identified transposable elements as drivers of genome size reduction during host adaptation.

## Results and Discussion

### Rhabdochlamydiae thrive in soil and freshwater environments

Previous analyses of metagenomic and 16S ribosomal RNA gene-based surveys predicted the Rhabdochlamydiaceae as one of the most diverse families within the phylum Chlamydiae [6, 10]. Since then, available sequencing data increased manifold, e.g., by one order of magnitude in the publicly available high throughput sequencing repository SRA, from ~1 000 TB in 2014 to ~10 000 TB in 2020 (trace.ncbi.nlm.nih.gov/Traces/sra/). To get an up-to-date overview we screened the SRA for 16S rRNA gene sequences using the database IMNGS [32]. This analysis suggested that the family Rhabdochlamydiaceae consists of at least 388 genus-level lineages and 14 051 species-level operational taxonomic units (OTUs; clustered using a sequence similarity threshold of 95 % and 99 %, respectively). We calculated this lower bound estimate using only sequences covering the V3-V4 region of the 16S rRNA gene as this is the most well-covered region in our dataset, comprising about 72 % of all sequences. Considering also other variable regions would likely result in an overestimation of OTUs as two sequences spanning different regions of the same 16S rRNA gene would appear as two separate genus-level OTUs in this analysis (see Materials and Methods). Compared to the few Rhabdochlamydiaceae full-length sequences reported to date, these estimates predict a staggering high natural diversity for members of this chlamydial family. A prime example lending support for this finding is a recent study investigating fecal microbiota from more than 400 insectivorous barn swallows during two breeding seasons [33]. The rhabdochlamydial 16S rRNA gene sequences detected in this longitudinal study alone contribute to 80 different genus-level lineages. The placement of representative sequences of all putative genus-level OTUs into a reference tree consisting of chlamydial full-length sequences illustrated the broad diversity of the Rhabdochlamydiaceae and showed that the predicted OTUs indeed span the entire family clade, including lineages both closely related and distant to previously recognized members (Figure 1).

**Figure 1:**
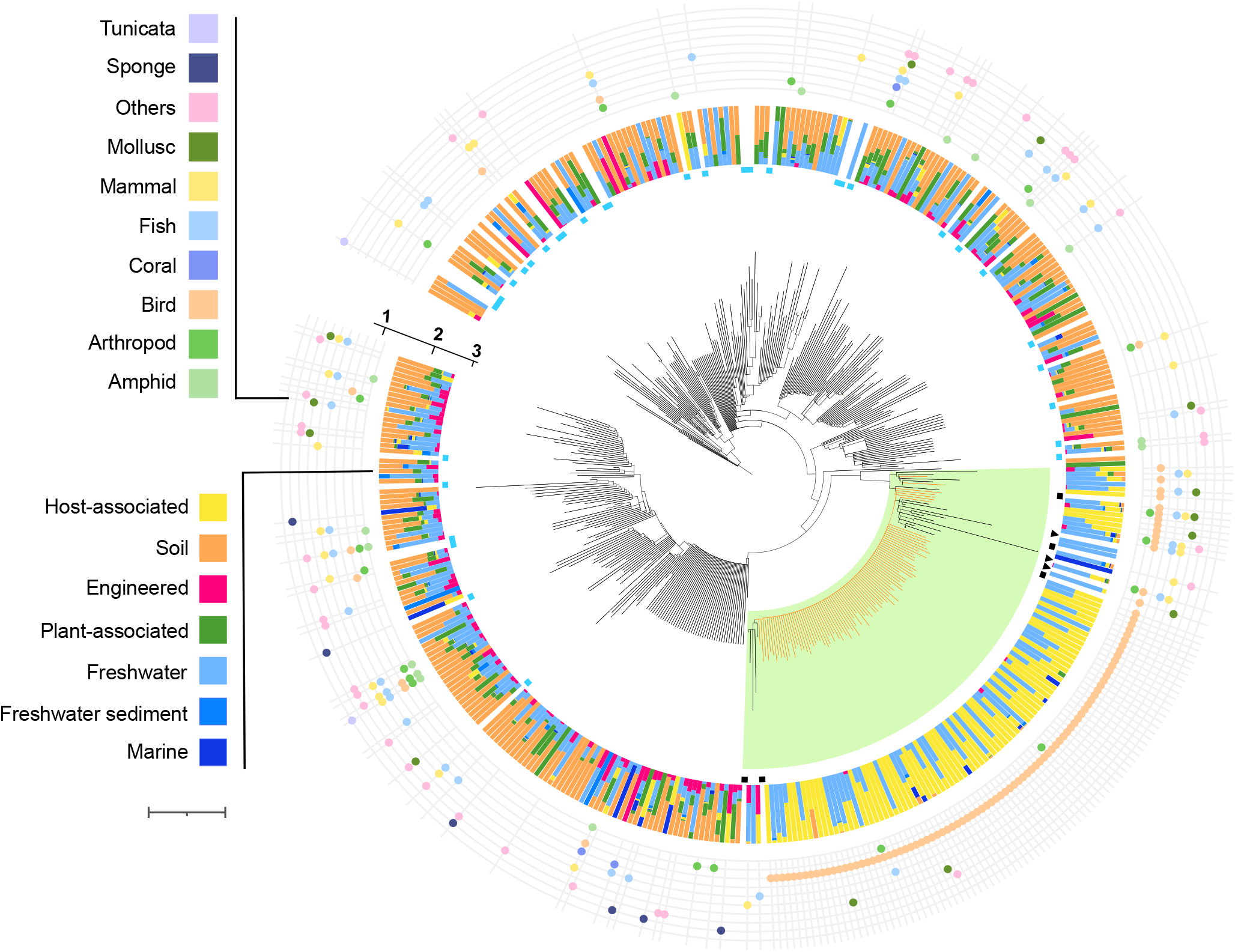
Rhabdochlamydiaceae are highly diverse and can be found in diverse environments and hosts. 16S rRNA gene tree of the family Rhabdochlamydiaceae, including full-length sequences (Supplementary figure S1) and partial sequences covering the V3-V4 region obtained from the IMNGS sequence database. For each of the 388 genus-level OTUs only one representative sequence (centroid) was included. The tree was rooted using other chlamydiae and members of the Planctomycetes-Verrucomicrobia-Chlamydiae (PVC) superphylum [36, 37] as an outgroup. Tree annotations from the outside to the inside: (**1**) represents the different host organisms for the category “Host-associated”, (**2**) indicates the relative abundance of environments where rhabdochlamydial 16S rRNA sequences were detected (**3**) represent full-length sequences (blue squares), full-length sequences associated with arthropod hosts (black squares), and 16S rRNA sequences of the genomes described in this study (black arrowheads). A monophyletic group including known arthropod-associated rhabdochlamydiae is highlighted in light green. Branches representing centroids including sequences from a longitudinal study of barn swallow feces are indicated in orange (Kreisinger et al. 2017), Scale bar indicates 0.1 substitutions per position in the alignment.

Although all known representatives of the family Rhabdochlamydiaceae are associated with arthropod hosts [25, 26, 28], our data show that most OTUs originate from soil (43 %) and freshwater (33 %) samples, suggesting the presence of additional, yet unknown hosts (Figure 1). Protists are abundant and important members of microbial communities in those environments [34, 35] and might thus serve as hosts for many of these lineages. Consistent with this, only 5 % of all identified rhabdochlamydial OTUs were detected in animal microbiomes from molluscs, birds, fish, and diverse mammals, and categorized as host-associated in our analysis (Figure 1). Most of these sequences, however, originate from feces or gut samples, and it is thus conceivable that rhabdochlamydiae are taken up with food and do not represent active infections. In fact, there is no general discernible pattern or pronounced correlation of phylogeny and relationship with environmental origins or putative host taxa in our dataset. We still noted a monophyletic group comprising all known arthropod associated Rhabdochlamydiaceae, i.e., the three described Rhabdochlamydia species. This clade contains in addition 107 genus-level lineages found in diverse environments, including many detected in feces from insectivorous birds (Figure 1). Taken together, our data suggests that while there is evidence for yet unknown environmental hosts, diverse animals may serve as either transient hosts or simply act as vectors for distributing rhabdochlamydiae through food uptake and excrements.

### Genome features and gene content distinguish rhabdochlamydiae from other chlamydial families

To learn more about members of the Rhabdochlamydiaceae, we next collected all available whole genome sequences and high quality MAGs (n=9; see Material and Methods; Table S1) and compared them to the most well-studied chlamydial families with known hosts, namely the Chlamydiaceae (human and other vertebrate hosts) [36] and the Parachlamydiaceae (amoeba hosts) [3]. We did not consider members of other described families due to the limited number of genome sequences and the lack of knowledge about their natural hosts, respectively. In addition, we determined the complete genome sequences of two rhabdochlamydiae from arthropod hosts: *R. porcellionis* infecting the woodlouse *Porcellio scaber* [25], and the new species *R. oedothoracis*, which is associated with the linyphiid dwarf spider *Oedothorax gibbosus* [27] (Table S2; for a formal candidatus species description see Text S1). In order to compare the different chlamydial families, we first clustered all genes into orthologous groups (OG) representing gene families [37, 38]. Next, we compared all genomes based on their gene content, i.e., abundance of gene families (Figure 2A, 2B). This analysis confirmed previous observations that the human and animal pathogens of the Chlamydiaceae clearly differ from the amoeba symbionts of the Parachlamydiaceae with respect to their genetic repertoire [2].Further, the number of genes shared within a chlamydial family is generally higher than the number of genes shared by the whole phylum [24]. These conserved family-specific genomic backbones have been interpreted to reflect adaptations to different niches or, as chlamydiae are obligate intracellular bacteria, to different hosts. Notably, our gene content analysis revealed that members of the Rhabdochlamydiaceae are clearly distinct from the Chlamydiaceae and the Parachlamydiaceae for both highly conserved (Figure 2A) and chlamydiae-specific gene families (Figure 2B). The different genome composition is also reflected in the degree of genome reduction, with rhabdochlamydiae showing intermediate genome sizes compared to the Chlamydiaceae and the Parachlamydiaceae (Figure 2C). Together, this suggests that the Rhabdochlamydiaceae have a different niche, for instance a different host range, in comparison to the other well-studied chlamydial families.

**Figure 2:**
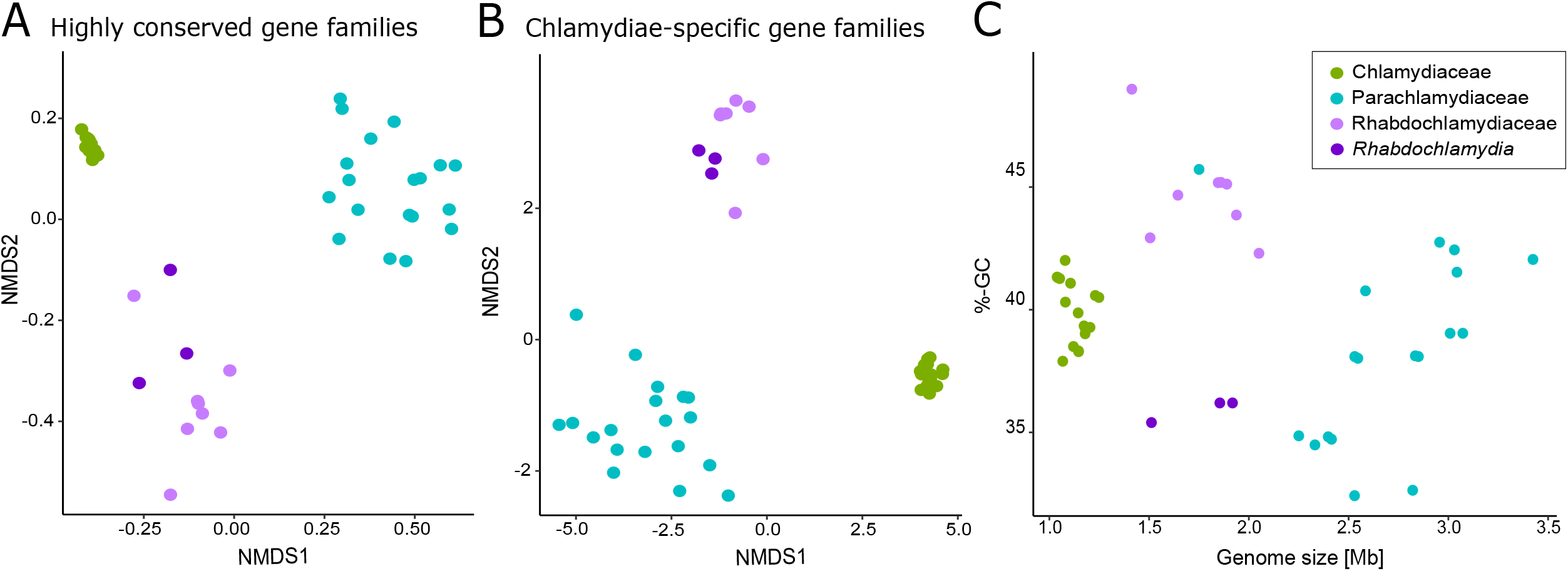
Rhabdochlamydiaceae genomes are distinct from those of vertebrate and amoeba-associated chlamydiae. Non-metric multidimensional scaling based on (**A**) highly conserved eggNOG OGs, and (**B**) chlamydiae-specific de novo clustered OGs of members of the families Rhabdochlamydiaceae, Parachlamydiaceae, and Chlamydiaceae. Each dot represents a genome, and the color represents the family. The genomes of known arthropod-associated rhabdochlamydiae are depicted in dark violet. The stress values indicate a good fit (A=0.06, B=0.07), (**C**) Correlation of genome size and GC-content for Parachlamydiaceae, Chlamydiaceae and Rhabdochlamydiaceae, respectively.

In many host-associated bacteria there is a correlation between genome size and GC content [39, 40], with smaller genomes tending to have a lower GC content. However, this does not seem to apply to chlamydiae [18, 40], suggesting evolutionary forces other than relaxed selection and genetic drift shaping the genomic GC content of members of this phylum. Several other factors are known to drive the base composition in bacteria including environmental conditions and niche-specialization [41, 42]. Within the family Rhabdochlamydiaceae we observe a clear divide with respect to the genomic GC content, with known arthropod-associated rhabdochlamydiae (i.e., the members of the genus *Rhabdochlamydia: R. helvetica*, *R. oedothoracis*, *R. porcellionis*) differing pronouncedly from those of other rhabdochlamydiae (35.4 %-36.2 % vs. 42.3 %-45.2 % on average; Figure 2C; Figure S2). This might indicate that the latter thrive in a different niche, i.e., are associated with hosts other than arthropods. However, more Rhabdochlamydiaceae genome sequences from arthropod hosts are needed to corroborate these observations.

### The Rhabdochlamydiaceae pangenome

To explore the genomic setup of the Rhabdochlamydiaceae in more detail we conducted a pangenome analysis. The pangenome describes all genes in a certain group of organisms and consists of genes present in all individuals in that group, the core genome, and genes that are specific for only some of them, referred to as accessory genome [43]. For this analysis we selected all nine Rhabdochlamydiaceae genomes from our dataset (Table S1; Figure S3). The family pangenome comprises 5 178 OGs of which 665 are present in > 90 % of all genomes, representing the core genome. This includes almost all of the genes constituting the chlamydial core genome [18, 24], such as the type III secretion system [44], nucleotide transport proteins (Ntt1/Ntt2) [45], the master regulator of the chlamydial developmental cycle (Euo)[46] as well as major effector proteins (CopN, Pkn5)[47, 48] that interfere with host cellular pathways. Further, glycogen metabolism is conserved among all Rhabdochlamydiaceae, this is consistent with the importance of glycogen as storage compound for many known chlamydiae [49].

The accessory genome contains lineage-specific genes representing adaptations to different niches [18, 24, 43, 50]. In general, the arthropod-associated Rhabdochlamydia species tend to have smaller accessory genomes (246-395 genes) than other members of the family Rhabdochlamydiaceae (366-588 genes) with unknown hosts (Wilcoxon rank sum test; p-value=0.05) (Figure S4). When we grouped the accessory genes into functional categories inferred from annotations in the eggNOG database, we could not recognize clear differences between the individual genomes (Figure S5). However, among the gene families differentiating known arthropod-associated rhabdochlamydiae from other Rhabdochlamydiaceae, i.e., those gene families that are unique to or completely missing in the genus Rhabdochlamydia, we found several genes associated with cell wall or membrane biosynthesis (Table S3). Whether any of these are related to the rod-shaped EBs and the characteristic five-layered cell envelope of arthropod-associated Rhabdochlamydia species remains to be determined [29, 30, 51].

### *The genus* Rhabdochlamydia

Next, we further focused on the genus *Rhabdochlamydia*, which represents the best studied clade in the family, because: (i) it includes the only cultured representatives of the Rhabdochlamydiaceae, (ii) the hosts of all three described species are known, and (iii) its members are represented by two closed and one high-quality draft genome, including one plasmid each. Calculation of the genome-wide average amino acid identity (AAI) confirmed their classification into a single genus (AAI >80 %; Figure S6; [52]). The *Rhabdochlamydia* genus pangenome comprises 1 875 OGs, where most of them belong to the core genome (1 007, 54%). The sizes of the accessory genomes vary between the species and correlate with genome size (Figure 3A). Between 21 % and 37 % of the accessory genomes mapped to known gene families in the eggNOG database; the larger proportion consists of orphan genes and genes with remote homology to genes of unknown function.

**Figure 3:**
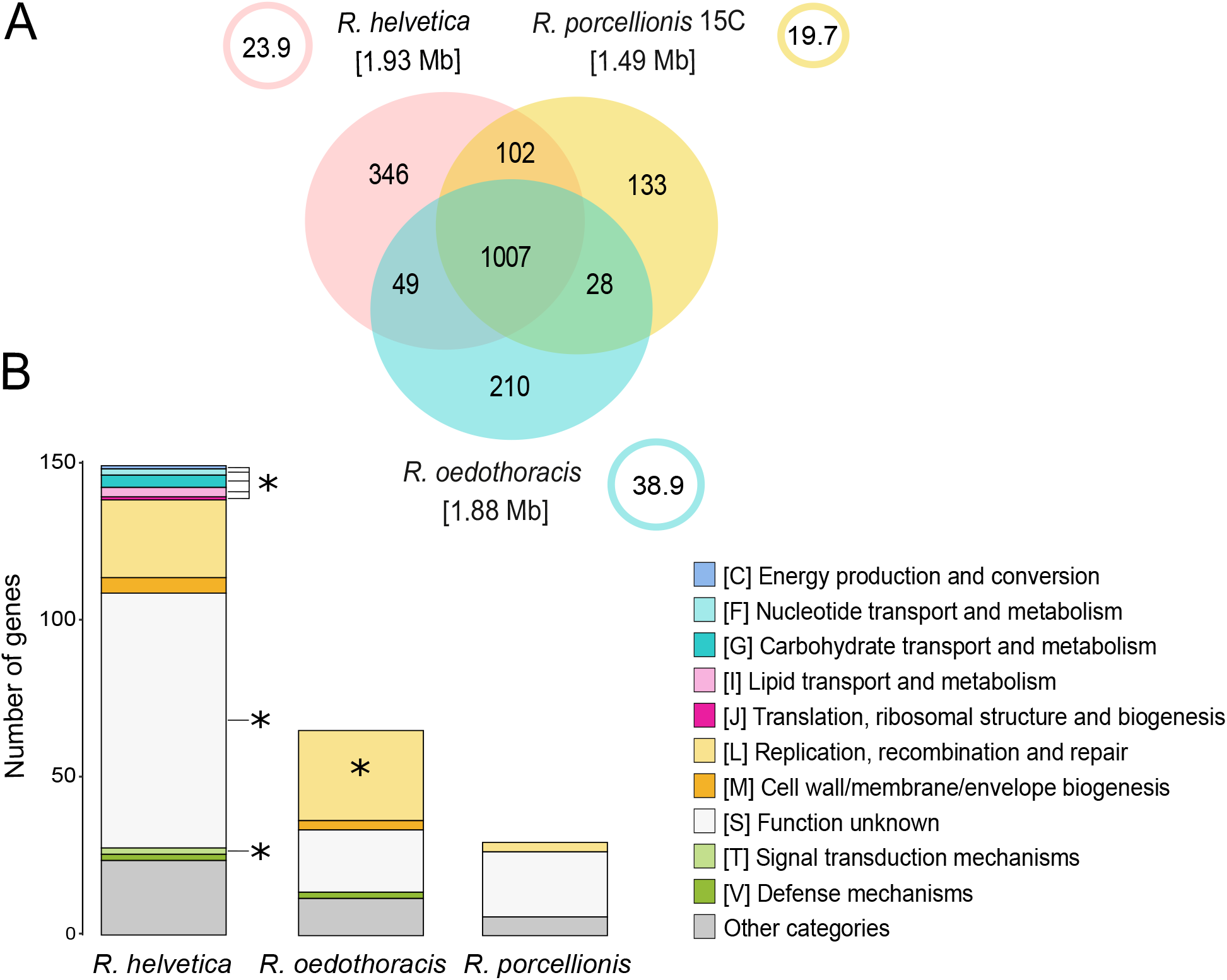
The pangenome of the genus Rhabdochlamydia. (A) Venn diagram representing the pangenome of the genus Rhabdochlamydia. The numbers represent the orthologous groups, i.e. gene families, shared between the respective genomes. The numbers in brackets represent the estimated genome sizes in Mb. The circles represent the plasmids, with plasmid sizes indicated in Kb. The large core genome reflects major metabolic pathways and other well-known conserved chlamydial features. (**B**) Stacked-bar chart showing functional categories of the subset of accessory genes mapped to eggNOG (only unambiguous annotations were used). Only selected categories are shown, the rest are combined into “Other categories”. Significantly enriched categories (false discovery rate adjusted p-value < 0.001) are marked with an asterisk. Differences in the accessory genomes reflect differences in host interaction and the degree of host adaptation.

We noted, however, that the genomes of *R. helvetica* and *R. porcellionis* include a complete pathway for the *de novo* synthesis of polyamines. Polyamines play an important role in virulence and response to various stressors [53–55]. The complete pathway is an unusual feature of chlamydial genomes [28], seems incomplete or absent in other rhabdochlamydial genomes, and is also absent in the closest cultured relative outside the Rhabdochlamydiaceae, *Simkania negevensis*.

All members of the genus *Rhabdochlamydia* carry a large plasmid between 20 and 39 kB in size (Figure 3A). Plasmids are small DNA molecules replicating independently from the chromosome, known to mediate horizontal gene transfer, and are considered important for the adaptation to different environments [56]. Plasmids have been identified as important drivers of genome evolution in the phylum Chlamydiae [57, 58], and the highly conserved Chlamydiaceae plasmid is recognized for its role in virulence in human and animal hosts [57–59]. In total, *Rhabdochlamydia* plasmids encode 83 proteins that belong to 39 different gene families. More than half of the gene families have representatives on at least one other *Rhabdochlamydia* chromosome or plasmid. This indicates a high degree of gene flow between chromosomes and plasmids, an observation also described for other chlamydial plasmids [57].

All *Rhabdochlamydia* plasmids contain genes considered to be important for plasmid maintenance in the *Chlamydiaceae*, such as Pgp2, plasmid partitioning protein ParA, and the integrase Pgp8 [59]. Interestingly, the *Rhabdochlamydia* plasmids encode major outer membrane (MOMP)-like proteins, in addition to the respective chromosomal copies. MOMPs are highly conserved among chlamydiae. They function as porins and adhesins and are prominently recognized by the host immune system in members of the Chlamydiaceae [60]. The MOMP-like proteins of *Rhabdochlamydia* show little to no similarity with the canonical MOMPs of the Chlamydiaceae. However, they belong to a large number of orthologs also found in the only distantly related chlamydiae *Waddlia chondrophila* and *S. negevensis*, with yet unknown function [18, 61, 62].

To more systematically compare the *Rhabdochlamydia* accessory gene sets, we performed an enrichment analysis taking into account functional category annotations from eggNOG (Figure 3B). The *R. oedothoracis* accessory genome is enriched in the category “replication, recombination and repair” (FDR adjusted p-value < 0.001), which includes transposases and genes for their maintenance. *R. helvetica* on the other hand is enriched in several categories and includes a large number of genes with unknown function (FDR adjusted p-value < 0.001) (Figure 3B). In addition, the accessory genomes of *R. oedothoracis* and *R. helvetica* include a range of functions that are linked to communication with the environment like defense mechanisms and cell wall/membrane/envelope biogenesis that are missing in *R. porcellionis*. Together with the smaller genome size of *R. porcellionis*, this indicates a prolonged association with the woodlouse host and may reflect an adaptation to the limited competition with other bacteria in the hepatopancreas - the target organ of infection [25, 63].

### Insertion sequences as key players in genome reduction

Reduced genomes are a hallmark of all chlamydiae [6, 18]. Yet, the evolutionary trajectories leading to their streamlined and highly specialized genomes are poorly understood. Members of the genus *Rhabdochlamydia* with their differences in genome size might offer an interesting perspective to learn more about the process of genome size reduction and host adaptation in these bacteria. To this end, one of the most striking differences between the known *Rhabdochlamydia* genomes is the presence of a high number of transposases in *R. oedothoracis* and its mere absence in the smallest genome of *R. porcellionis*.

Transposases are indicative of transposable elements (TEs), which in their simplest version as insertion sequences (ISs) contain only a transposase gene flanked by inverted repeats [64]. There are several reports of ISs being associated with genome reduction in beneficial bacterial symbionts [65–71]. In most of these cases, the symbionts were recently acquired from a free-living stage. During the adaptation to the host and the intracellular environment, the symbionts accumulate ISs in their genomes [72]. The ISs may interrupt genes that then accumulate mutations, especially deletions, as a consequence of relaxed selection, which ultimately leads to a reduction of genome size [72]. Here, we suggest that a similar process drove the evolution of *Rhabdochlamydia* genomes.

As ISs are known to cause breaks in genome assemblies and are often not properly annotated by automated tools [73], we limited our in-depth analysis to the closed genomes of *R. oedothoracis* and *R. porcellionis*, and we manually curated transposase annotations (Figure 4B; see methods for details). In total, we could identify 415 transposase genes in *R. oedothoracis* and only 19 in *R. porcellionis*. Apart from 129 transposases in *R. oedothoracis*, most of those do not appear to be functional; they are either truncated, contain premature stop codons, or are interrupted by other transposases (Table S4). Notably, (functional) transposases are also encoded on the plasmid of *R. oedothoracis* yet absent on other rhabdochlamydial plasmids (Table S4). It was shown previously that plasmids need to exceed a certain minimum size (~20 kB) to be able to carry TEs [64]. This threshold would explain the absence of transposases on the plasmids of *R. porcellionis*. The presence of representatives of the most abundant chromosomal transposase families on the plasmid of *R. oedothoracis*, however, may suggest a role of the plasmid in IS expansion. The higher copy number of plasmids and their replication independent of the chromosome [56] might support the proliferation of ISs.

**Figure 4:**
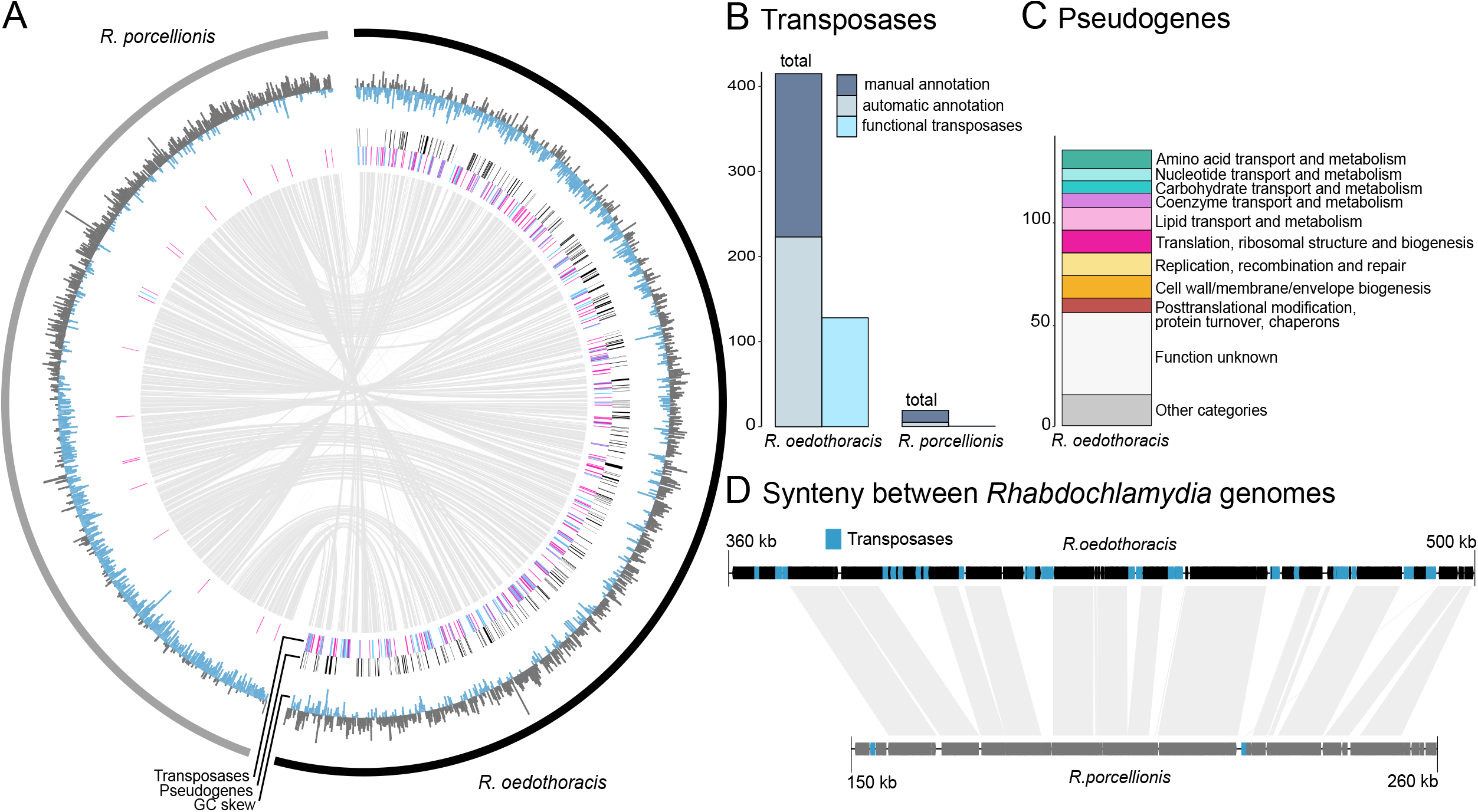
Transposable elements and pseudogenization in the genus Rhabdochlamydia. (**A**) Comparison of the R. oedothoracis (black) and R. porcellionis (grey) genomes. The outermost ring represents the GC skew. The second ring shows the positions of predicted pseudogenes indicated by black lines. Transposases are shown in blue (functional) or orange (degraded) in the third ring. The center of the plot shows syntenic regions between the two genomes based on blastn (eval < 0.001). Transposases are spread throughout the genome of R. oedothoracis causing breaks in the synteny to R. porcellionis. (**B**) Number of manually and automatically annotated transposases in the closed genomes of R. oedothoracis and R. porcellionis. Manual annotation and curation are essential to comprehensively identify functional and degraded transposase genes. The fraction of functional transposases is depicted in light blue. (**C**) Functional categories of predicted pseudogenes with an eggNOG annotation in R. oedothoracis (only unambiguous annotations were used). Only the most abundant categories (n < 5) are shown, the rest are combined to “Other categories”. (**D**) Alignment of a selected syntenic genome region of R. oedothoracis (black) and R. porcellionis (grey) illustrating the role of ISs in genome size reduction. Transposases are shown in blue, genes are represented by black and grey boxes, respectively.

Apart from a high number of TEs, increased pseudogenization is indicative for genomes under degradation [72]. We therefore used pseudofinder [74] to identify genes under relaxed selection in the genome of *R. oedothoracis* by comparing it to *R. porcellionis*. This approach assumes that due to its small size and the low number of transposases, most genes are under purifying selection in the reference genome of *R. porcellionis*. In total, 276 *R. oedothoracis* genes were marked as cryptic pseudogenes i.e., genes that are structurally intact but likely experience relaxed selection (dN/dS ratios >= 0.3) [68]. A broad range of functions is affected by this ongoing pseudogenization, including diverse metabolic pathways, as well as genes involved in replication and regulation (Figure 4C).

Taken together, with its small size and the low number of transposases, the genome of *R. porcellionis* is the most streamlined genome in the genus, suggesting an ancient association with its *P. scaber* host. In contrast, there is still evidence for the process of genome reduction in the case of *R. oedothoracis*, given the high number of (functional) transposases and pseudogenes, possibly as a consequence of a relatively recent host switch. Notably, the distribution of transposases in the *R. oedothoracis* genome correlates with positions where the synteny of the two genomes is disrupted (Figure 4A, 4D). This further illustrates the putative role of ISs in genome rearrangements and genome size reduction in *Rhabdochlamydia*. Consistent with this, there is further evidence for a nascent stage of genome reduction in *R. oedothoracis*: Although the GC content of the rearrangement regions generally matches that of the surrounding regions, the characteristic asymmetrical pattern of circular chromosomes in cumulative GC skew analyses [75] is less pronounced (Figure 4A).

To learn more about the origin of the transposases present in the genome of *R. oedothoracis*, we performed phylogenetic analyses for the three most abundant transposase families with functional representatives in *R. oedothoracis*. Surprisingly, all investigated transposases showed a phylogenetic relatedness to transposases found in other chlamydiae (Figure S7), suggesting the existence of an ancient pool of transposases in chlamydial ancestors and sequential loss in several lineages.

### *A scenario for the evolution of the genus* Rhabdochlamydia

Our observations regarding diversity, environmental distribution, and genomics of members of the family Rhabdochlamydiaceae provide clues about genome evolution and the adaptation of chlamydiae from symbionts of unicellular eukaryotes to animal hosts.

We show that members of the Rhabdochlamydiaceae are highly diverse, occur in different environments and mostly lack a clear association with animal hosts (Figure 1). This suggests that the majority of rhabdochlamydiae infect other, yet unknown and likely unicellular hosts. Surprisingly, however, members of the Rhabdochlamydiaceae differ pronouncedly in their genetic make-up and genome size from recognized chlamydial symbionts of heterotrophic amoeba (Figure 2A, 2B, 2C). Yet, there is a wide range of protists with very different lifestyles e.g., phototrophic, or saprotrophic protists feeding on decaying organic matter, that could serve as natural hosts for rhabdochlamydiae [34]. According to the “melting pot” hypothesis, symbionts in amoebae retain larger genomes than closely related bacteria infecting animals as there is a high level of competition and possibilities for gene acquisition by lateral gene transfer in amoebae that feed on complex microbial communities [76, 77]. In phototrophic or saprotrophic protists, the competition and interaction with other bacteria would be much lower, leading to smaller genome sizes and differences in the genetic repertoire as seen for the Rhabdochlamydiaceae (Figure 2A, 2B, 2C). We thus suggest that members of the family include widespread symbionts of protist hosts different from the phagotrophic, free-living amoeba recognized so far as hosts for other chlamydiae. Of note, there is recent evidence for diverse chlamydial symbionts including rhabdochlamydiae in the cellular slime mold *Dictyostelium discoideum* [78].

Within the family Rhabdochlamydiaceae, the similar GC content, a large core genome and shared membrane features distinguish the genus *Rhabdochlamydia* from all other members (Figure 2A, 2C). This is consistent with them sharing a similar niche in arthropod hosts and putatively originating from rhabdochlamydiae thriving in environmental protists (Figure 1, 3). By infecting hosts equipped with an innate immune response, members of the genus *Rhabdochlamydia* might represent an intermediate step towards adaptation of chlamydiae to vertebrate animals with adaptive immunity. In this scenario, food or water would be a conceivable entry route for the uptake of protist-associated rhabdochlamydiae by arthropod hosts. We suggest that the subsequent transition and adaptation to arthropod hosts triggered genomic changes in the last common ancestor of *Rhabdochlamydia* species, resulting in reduced and specialized genomes of extant members of the genus. This process was putatively facilitated by IS expansion, inactivating genes under relaxed selection and eventually leading to genome size reduction (Figure 5). Genome reduction mediated by transposable elements is common in inherited, vertically transmitted beneficial symbionts [41, 67]. To our knowledge, such a scenario has not yet been described for horizontally transmitted intracellular bacteria representing commensals or pathogens as it is the case for members of the phylum Chlamydiae. The extent of genome streamlining might be dependent on the arthropod hosts, the site of infection and the extent of competition with other microbes. The digestive glands of *P. scaber*, the target organ of *R. porcellionis*, for example, harbors only a few other bacteria [63]. The same is true for the hindgut of the spider host of *R. oedothoracis* [27]. The tick *Ixodes ricinus*, on the other hand, contains a diverse microbiome, creating a more competitive environment for *R. helvetica* and opportunities for genetic exchange [79].

**Figure 5:**
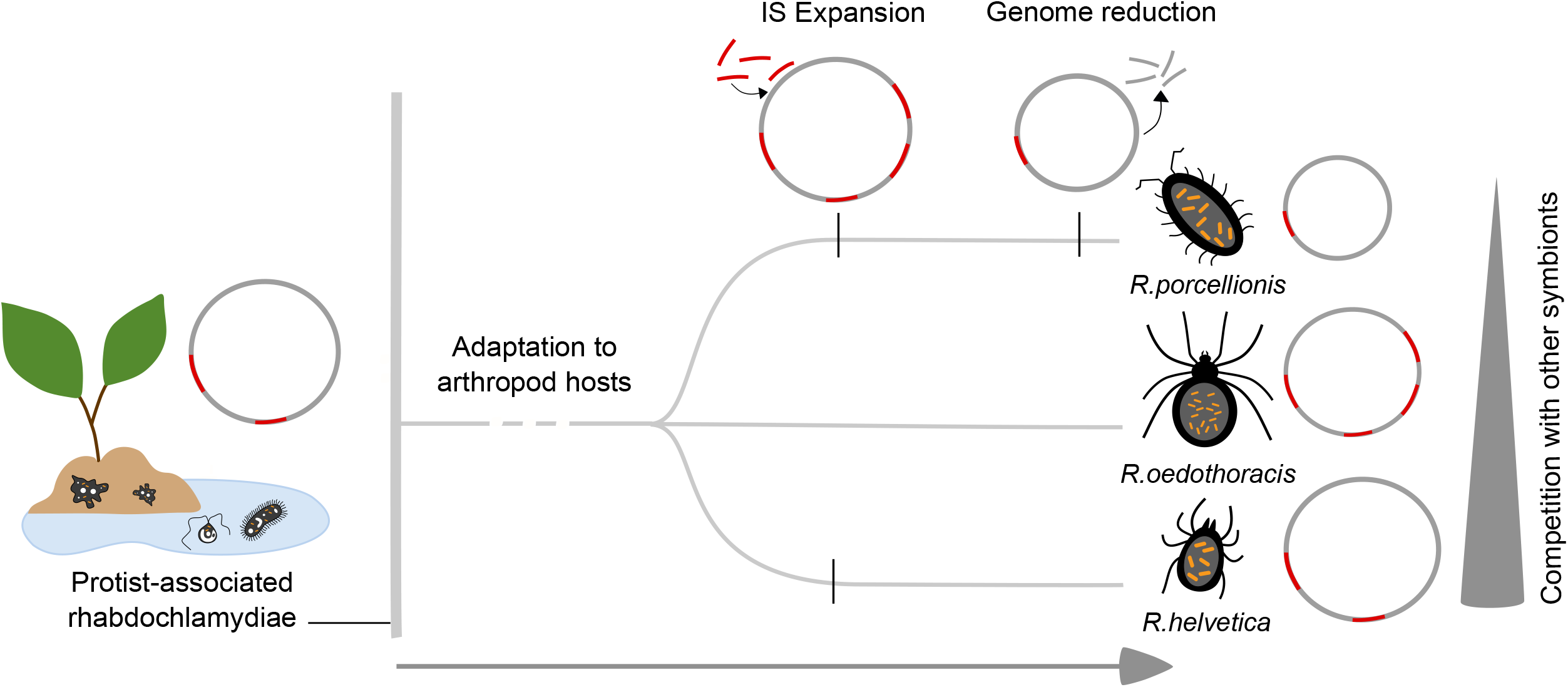
Genome evolution and adaptation of Rhabdochlamydia species to arthropod hosts. A scenario for the evolution of arthropod-associated rhabdochlamydiae: While most rhabdochlamydiae are putatively associated with protist hosts, a small monophyletic group established that is able to infect arthropods. The adaptation to arthropod hosts was possibly facilitated by expansion of ISs and subsequent genome streamlining, impacted by competition with other bacteria and conditions in the new host.

In conclusion, we have demonstrated that Rhabdochlamydiaceae are distributed globally and represent a major, yet heavily underexplored chlamydial group. We show that they provide opportunities to study adaptation and genome evolution of chlamydiae during the transition from protist to animal hosts. We have identified transposable elements as an important factor underlying genome size reduction in the phylum Chlamydiae, and we propose a scenario for the adaptation of *Rhabdochlamydia* species to their arthropod hosts. A limitation of our study is the low number of available high-quality *Rhabdochlamydia* genome sequences. Sequencing more arthropod-associated chlamydiae is needed to verify the evolutionary scenario proposed here. Further, the in-depth analysis of members of the family Rhabdochlamydiaceae is hampered by the dramatic lack of cultured representatives and information about host organisms. Future efforts targeting understudied protist taxa and recovering symbionts together with their hosts from complex environmental samples might help to overcome this. Taken together, the current study provides a comprehensive framework for investigating the ecology and evolution of one of the most widespread lineages within the phylum Chlamydiae.

## Materials and Methods

### 16S rRNA gene phylogeny

We downloaded all available unique near-full length 16S rRNA gene sequences of chlamydiae (n=233) and other Planctomycetes-Verrucomicrobia-Chlamydiae (PVC) members (n=205) from SILVA v138 SSU Ref NR 99 [80] and added 78 16S rRNA genes from published chlamydial genomes from RefSeq [81] and GenBank [82]. In addition, we added 79 near-full length chlamydial sequences from Schulz et al. 2017. We dereplicated the sequences at 99 %-identity using USEARCH (v11) [84] with “-cluster_smallmem” and aligned the clustered sequences with SINA [85]. Afterwards, we trimmed the alignment with trimAl (v1.4.15) [86] “-noallgaps” and removed the highly variable positions using noisy (v1.5.12) [87]. The phylogenetic tree was then calculated with IQ-TREE (v1.6.2) [88]. Model testing was performed with “-m TESTNEW” (Best model: SYM+R9), and initial support values were inferred from 1 000 non-parametric bootstraps using “-bb 1000”. The final tree was edited and visualized using iTOL (v4) [89].

### 16S rRNA gene-based diversity and environmental distribution

We queried the IMNGS database, which is a collection of pre-clustered NCBI SRA sequencing data [32] on 09 June 2020 for 16S rRNA genes with at least 90 % identity to the reference 16S rRNA sequence of *R. porcellionis* 15C. We removed singletons, only kept sequences >400 bp, and removed duplicates and sequences with ambiguous bases using mothur (v.1.42.3). 16S rRNA genes were aligned to SILVA Ref NR 99 SSU (v138) [80] using mothur (v.1.42.3)[90], and the alignment was trimmed with trimAl (v1.4.15) [86] using the “ noallgaps” parameter. Afterwards, we clustered the sequences in OTUs using USEARCH (v11.0.667) [84] “-cluster_otus” to reduce redundancy, and finally on 95 % sequence identity level using “-cluster_smallmem”. In order to belong to one cluster, sequences were required to overlap to 90 % (“-query_cov 0.9”). Centroid sequences were aligned to the 16S rRNA full-length alignment using MAFFT (v7.427) (“--addfragments”) [91], and variable positions were removed using trimAl (“-selectcols”) (v1.4.15) [86]. We then placed the centroids to the 16S rRNA full-length reference tree using EPA-ng (v0.2.1)[92] (model: SYM+R9), and manually selected all centroids that were placed in the family Rhabdochlamydiaceae. This step significantly reduced the number of centroids from 2 162 to 938. For the final tree, only rhabdochlamydiae centroids were placed into the 16S rRNA full-length tree. We selected only sequences covering the V3-V4 region of the 16S rRNA gene as considering also other variable regions would likely result in an overestimation of OTUs as two sequences spanning different regions of the same 16S rRNA gene would appear as two separate genus-level OTUs in this analysis. When considering also those sequences covering other 16S rRNA gene regions, we retrieved an additional 550 genus-level OTU candidates (262 OTUs for V4-V5; 87 for V5-V6; 201 for V6-V8). The final tree was edited and visualized using iTOL (v4) [89]. For the analysis of the relative abundance in the environment of rhabdochlamydiae centroids in total 14,051 sequences were analyzed. The metadata was provided by IMNGS and retrieved from the SRA. The broad categories provided by the SRA were manually curated and each rhabdochlamydiae sequence assigned to one of the following categories: freshwater, freshwater-sediment, marine, plant-associated, soil, and host-associated. The sequences assigned to host-associated were further categorized based on the organisms they originated from. Sequences that originated from gut or stool samples were also classified as host associated. In total, 670 sequences were assigned to the category host-associated, 4 515 to freshwater, 141 to freshwater-sediment, 6 002 to soil, 1 714 to plant-associated, 194 to marine and 815 to engineered. The bar charts were created by counting the total number of sequences represented by a centroid and calculating the relative abundances for each category.

### *Genome sequencing and assembly* - R. porcellionis *15C*

*R. porcellionis* 15C was cultivated in Sf9 insect cells (*Spodoptera frugiperda*) as described in Sixt et al. 2013. For DNA isolation we harvested Sf9 cells infected with *R. porcellionis* 15C and lysed the host cells with lysis buffer (1M Tris-HCl, 0.5 M EDTA, 5 M NaCl, SDS, Proteinase K). Afterwards, the host DNA was digested using DNase I (1 U/μL, Thermo Scientific; Thermo Fisher Scientific; Waltham, MA, USA). Bacterial gDNA isolation was carried out using the DNeasy Blood and Tissue Kit (Qiagen; Hilden, Germany). To remove remaining RNA, we treated the isolated gDNA with RNAse A (10 mg/mL, Thermo Scientific; Thermo Fisher Scientific; Waltham, MA, USA). Finally, we checked the quality of the gDNA using Qubit4 (Invitrogen; Thermo Fisher Scientific; Waltham, MA, USA) and the dsDNA HS Assay Kit (Invitrogen; Thermo Fisher Scientific; Waltham, MA, USA) and Nanodrop 1000 Spectrophotometer (Thermo Fisher Scientific; Waltham, MA, USA).

Before library preparation for the long read sequencing the gDNA was measured with Nanodrop and the length of the DNA fragments was measured with a Bioanalyzer. Library preparation was done using the Ligation Sequencing Kit (Oxford Nanopore, Oxford, UK; ONT). Sequencing was performed on an Illumina HiSeq 2000 platform (Illumina, San Diego, CA, USA), using the 100-bp-paired-end sequencing mode. Additional long-read sequencing was performed using a MinION sequencer (Oxford Nanopore, Oxford, UK).

For the assembly we trimmed the Illumina reads using bbduk (v37.61) (sourceforge.net/projects/bbmap/) (“-qtrim=rl-trimq=18-minlen=70”) and removed adapters and barcodes from the Nanopore reads using ONT’s qcat (“--trim”). We assembled the Illumina and Nanopore reads in a hybrid assembly using unicycler (v0.4.6) [94]. The quality of the assembly was checked by visually inspecting the assembly graph [95] and checkM (v1.0.18)[96].

### *Genome sequencing and assembly* - R. oedothoracis *W744*

DNA was isolated from a single field-captured *O. gibbosus* individual from the Walenbos population (W815). DNA isolation and Illumina sequencing were carried out as described in Hendrickx et al. 2021. The Illumina assembly was done using SPades (v3.9.1, “--meta”) [97]. The contigs were then binned using mmgenome [98]. Finally, reads were mapped to the metagenome assembled genomes (MAGs) and reassembled with SPades (v3.9.1, “--meta”) [97]. The quality of the MAGs was checked with checkM (v1.0.6) [96].

PacBio sequencing data from *O. gibbosus* individual W744 (Walenbos population) were obtained from [99]. PacBio reads were classified using a custom Kraken (v2.0.8) database including reference libraries for archaea, bacteria, viruses, protists, humans, fungi, and plants as well as MAG W815, and all reads classified as Rhabdochlamydiaceae were collected (MAG W744) [100].

PacBio reads were mapped to MAG W815 and W744, respectively using minimap2 (v2.17) [101]. As the coverage of the PacBio data was too high, the mapped reads were subsampled to a coverage of 70x. Finally, the reads mapped to MAG W815 and W744 were merged, and duplicates were removed. Illumina reads were mapped to MAG W815 and W744 using bbmap (v37.61) (sourceforge.net/projects/bbmap/) and merged and deduplicated afterwards. The final sets of Illumina and PacBio reads were then used for a hybrid assembly using unicycler (v0.4.6) [94]. The quality of the assembly was checked by visually inspecting the assembly graph [95] and checkM (v1.0.18) [96].

### Dataset compilation, quality control, and annotation

We downloaded 36 chlamydial reference genomes from GenBank/ENA/DDBJ [82] and RefSeq [81] on 25 June 2019 and added nine high-quality MAGs from the Genomes of the Earth’s Microbiome initiative [102]. Only genomes with a completeness >94 % and containing neither detectable strain heterogeneity nor contaminations were used, resulting in nine genome sequences and MAGs from the Rhabdochlamydiaceae, 17 Chlamydiaceae, and 19 Parachlamydiaceae genomes (Table S1). The quality of the genomes was checked using checkM (v1.1.3, “‘taxonomy_wf domain Bacteria”) [96], and basic statistics were calculated using QUAST (v5.0.2)[103]. Initial gene calling and annotation was performed with prokka (v1.14.6,” --mincontiglen 200”, “--gram neg”) [104].

The assembled genomes from *R. porcellionis* 15C and *R. oedothoracis* were annotated using prokka (v1.14.6) [104]. In addition, RNAs were annotated using the Rfam database [105] and cmscan (v1.1.3, “--cut_tc”, “--mid”) [106] and tRNAscan-SE (v2.0.5) [107]. The origin of replication was determined using the OriginX (v1.0) software [108]. Transposases were manually annotated by searching transposase sequences predicted by prokka against the ISfinder database [109] and manually curating the annotations using UGENE [110]. The *R. helvetica* genome contained in total 41 transposases predicted by prokka. This genome could, however, not be manually curated as it is not complete and thus neither the absence nor the misassembly of transposases can be excluded.

### Pangenome analysis

We mapped all proteins against the eggNOG database (v4.5.1) [37] using emapper (v1.0.1, “-d bact”) [111] to cluster them into orthologous groups. For all unmapped proteins we performed an all-against-all blastp search and clustered proteins with an e-value < 0.001 *de novo* with SiLiX (v1.2.11) with default parameters [38]. We used the following definitions for the pangenome components: core - present in more than 90% of genomes; accessory - present in only one of the genomes. Only for the pangenome of the genus *Rhabdochlamydia* we required a core protein to be present in all the genomes. For functional annotation we used eggNOG (v4.5.1) [37], and blastp [112] against the NCBI nr database for the *de novo* OGs. Further, we mapped all proteins to the Kyoto Encyclopedia of Genes and Genomes (KEGG) [113] orthologs (KOs) using GhostKOALA (v2.2) [114].

### *Comparison of* R. oedothoracis *and* R. porcellionis *genomes*

We used pseudofinder (v1.0) [74] and the “selection” function to identify genes under degradation in the genome of *R. oedothoracis* W744 in comparison to *R. porcellionis* 15C. Pseudofinder identifies homologous sequences in the two genomes and calculates the ratio of non-synonymous to synonymous substitution rates (dN/dS) for each set of genes. We used a threshold of 0.3 to distinguish between pseudogenes (> 0.3) and genes under purifying selection (<= 0.3).

To show synteny between *R. porcellionis* and *R. oedothoracis* the two genomes were blasted against each other using blastn [112]. Further, GC skews were calculated for both genomes using a custom python script (window size= 1000). The genomes were visualized using Circos (v0.69.9) [115]. To show disruption of synteny by transposases in more detail a short syntenic segment (*R. oedothoracis:* 360-500 kb, *R. porcellionis:* 150-260 kb) was picked and visualized using the “genoplotR” package (v0.8.11) [116] in R (v4.0.3) [117].

### Statistical analysis

All statistical tests and data analysis were performed in R (v4.0.3) [117] and visualized using ggplot2 (v3.3.3) [118]. NMDS was calculated using eggNOG (v 4.5.1) and *de novo* clustered OGs and the “metaMDS ^11^ function (“vegan” package v2.5-7)[119] using “distance=bray’”. To test whether members of the genus *Rhabdochlamydia* are associated with smaller accessory genomes we calculated the size of accessory genomes for all nine Rhabdochlamydiaceae genomes and used the “wilcox.test” function (“stats’’ package v4.0.3) for statistical evaluation. The enrichment analysis of functional categories based on eggnog (v 4.5.1) was carried out using a hypergeometric test with the “phyper” function (“stats’’ package v4.0.3). The p-value was corrected using the “p.adjust” function (“stats’’ package v4.0.3) and “method = BH”. We considered p-values < 0.001 as significant.

## Data availability

16S rRNA gene data used in this study are available via the SILVA database (https://www.arb-silva.de/) and IMNGS database (https://www.imngs.org/). Metadata for sequences received from the IMNGS database can be accessed via the Sequence Read Archive (SRA, https://www.ncbi.nlm.nih.gov/sra). Genome sequences generated in this study have been deposited in GenBank under the accession numbers CP075585-CP075586 (*R. porcellionis*) and CP075587-CP075588 (*R. oedothoracis*). Accession numbers for reference genomes and Metagenome-assembled genomes (MAG) are available in Supplementary Table S1. Metagenomic data are available through the IMG/M portal (https://img.jgi.doe.gov/). MAG sequences from the Genomes from Earth’s Microbiomes initiative are available at https://genome.jgi.doe.gov/GEMs. The collection of genomes and proteomes and the data of the IMNGS search used in this study are available at zenodo (https://10.5281/zenodo.4723235).

## Acknowledgement

Library preparation and long read sequencing was performed by the Next Generation Sequencing Facility at Vienna BioCenter Core Facilities (VBCF), member of the Vienna BioCenter (VBC), Austria. Illumina sequencing was carried out by the Norwegian High-Throughput Sequencing Centre (NSC), Oslo. The Life Science Compute Cluster (LiSC; http://cube.univie.ac.at/lisc) was used for computational analysis.

