## Supplementary Information for "Ecology and evolution of chlamydial symbionts of arthropods"

**Supplementary Text S1**: Description of *'Candidatus* Rhabdochlamydia oedothoracis'

[Rhab.do.chla.my'di.a. Gr. fem. n. rhabdos stick, rod; N.L. fem. n. Chlamydia taxonomic name of a bacterial genus; N.L. fem. n. Rhabdochlamydia rod-shaped chlamydiae, referring to the rod-like shape of elementary bodies; oe.do.tho.ra.cis. N.L. gen. n. oedothoracis of Oedothorax, pertaining to the taxonomic genus name of the host organism (a dwarf spider)].

*'Candidatus* Rhabdochlamydia oedothoracis' infects the spider *Oedothorax gibbosus* [1] and is yet uncultured; its genome sequence was reconstructed from a whole genome sequencing approach of the spider host [2]. Based on its 16S rRNA gene sequence, genomic average nucleotide identities (ANI), and its phylogenetic placement in 16S rRNA and marker gene-based analyses, *'Candidatus* Rhabdochlamydia oedothoracis' (genome sequence accession no. CP075588) belongs to the genus *Rhabdochlamydia* in the family Rhabdochlamydiaceae within the phylum Chlamydiae. *'Candidatus* Rhabdochlamydia oedothoracis' shows a 16S rRNA gene sequence identity of 98 % to *R. porcellionis* (accession no. CP075586) and 98 % to *'Candidatus* Rhabdochlamydia helvetica' (accession no. PRJEB24578). Further, the ANI of *'Candidatus* Rhabdochlamydia oedothoracis' and *'Candidatus* Rhabdochlamydia helvetica' amounts to 88.40 %, and the ANI of *'Candidatus* Rhabdochlamydia oedothoracis' and *R. porcellionis* is 89.85 %.

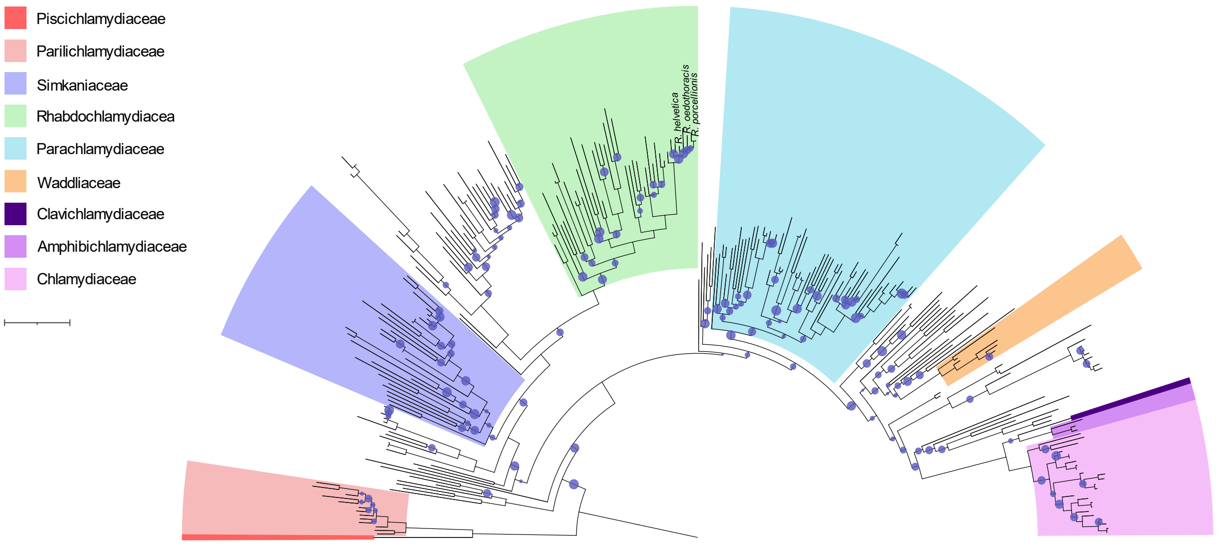


**Supplementary figure 1:** Full-length 16S tree of the phylum Chlamydiae. Tree is based on full-length alignment (99 % centroids) of 16S sequences retrieved from SILVA, RefSeq, Genbank and Schulz et al. (2017) [3-6]. Tree was rooted using other members of the PVC phylum as an outgroup. Rhabdochlamydiaceae are shown in green. Bootstrap values are depicted by circle sizes (0-0.8). Tree scale: 0.1.

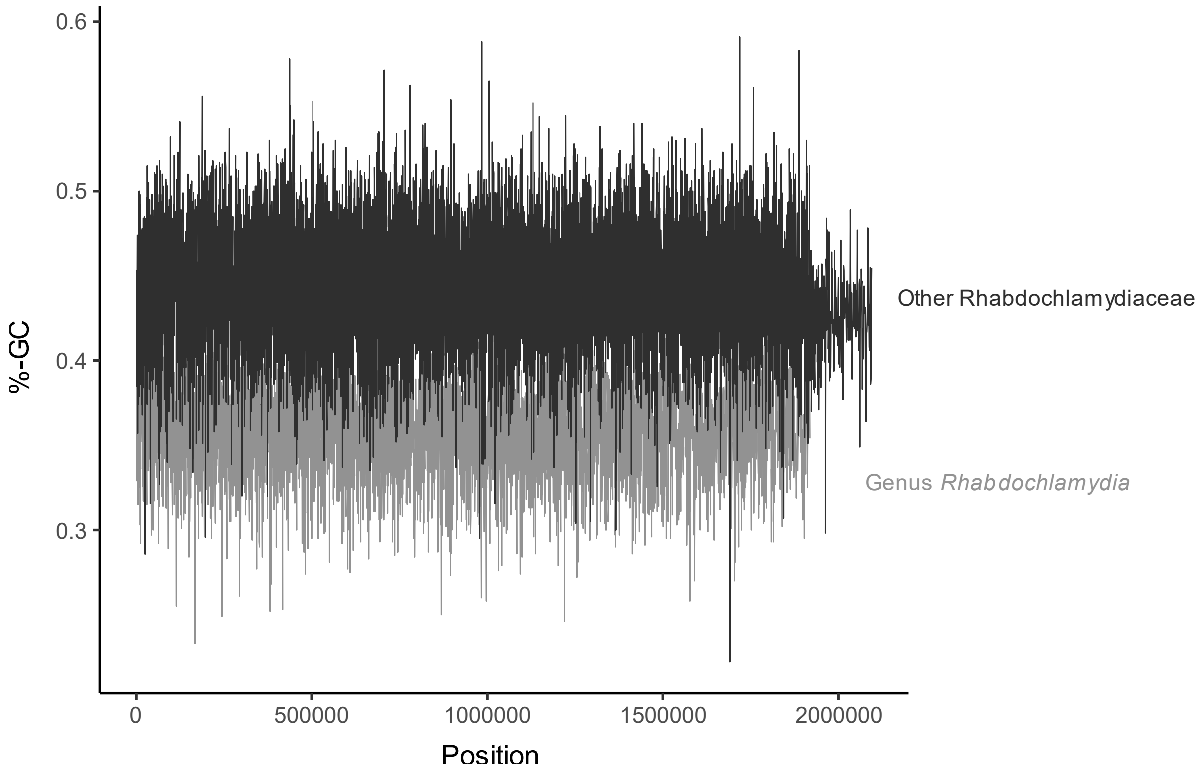


**Supplementary figure 2:** GC content (window size= 1,000) along rhabdochlamydiae genomes. CG content was calculated using a custom python script. Genomes belonging to the genus *Rhabdochlamydia* (*R. porcellionis*, *R. oedothoracis*, *R. helvetica*) are shown in light grey and three selected MAGs belonging to the family Rhabdochlamydiaceae but not to the genus *Rhabdochlamydia* (1095360-24, 1062783-10, 3300005529-103) are shown in dark grey.

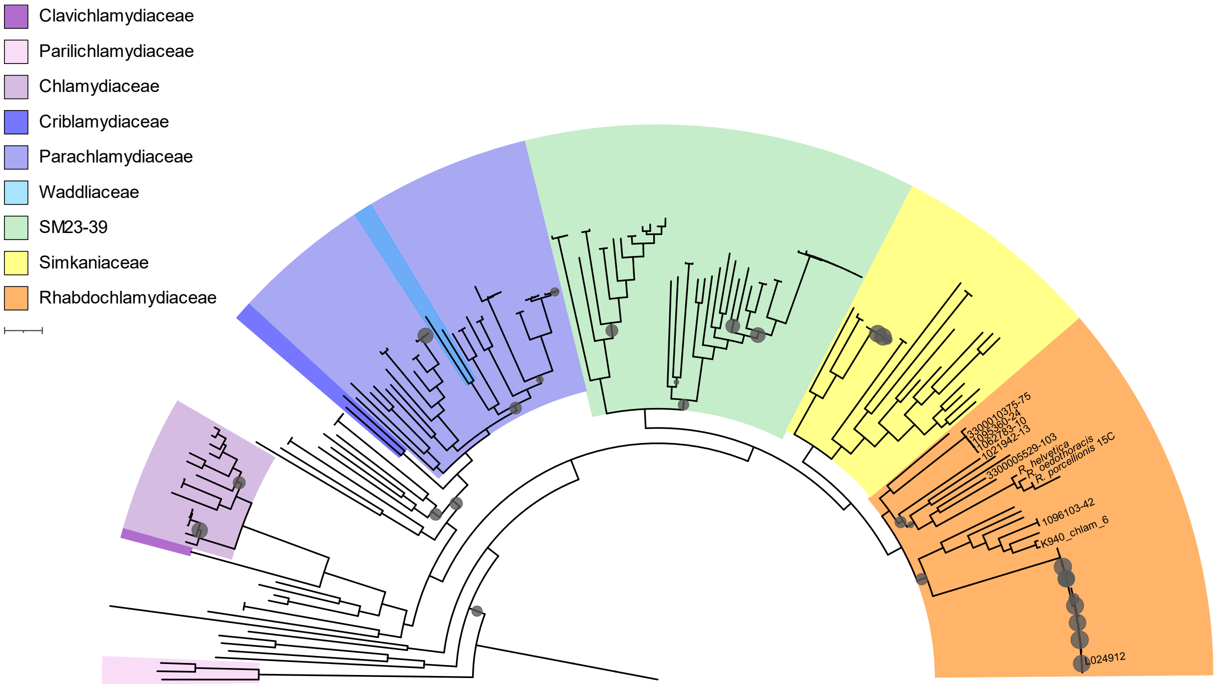


**Supplementary figure 3:** Marker-gene tree of the phylum Chlamydiae. Rhabdochlamydiaceae are shown in orange. Genomes used in this study are labeled accordingly. Bootstrap values are depicted by circle sizes (0-0.8). Tree scale: 0.1.

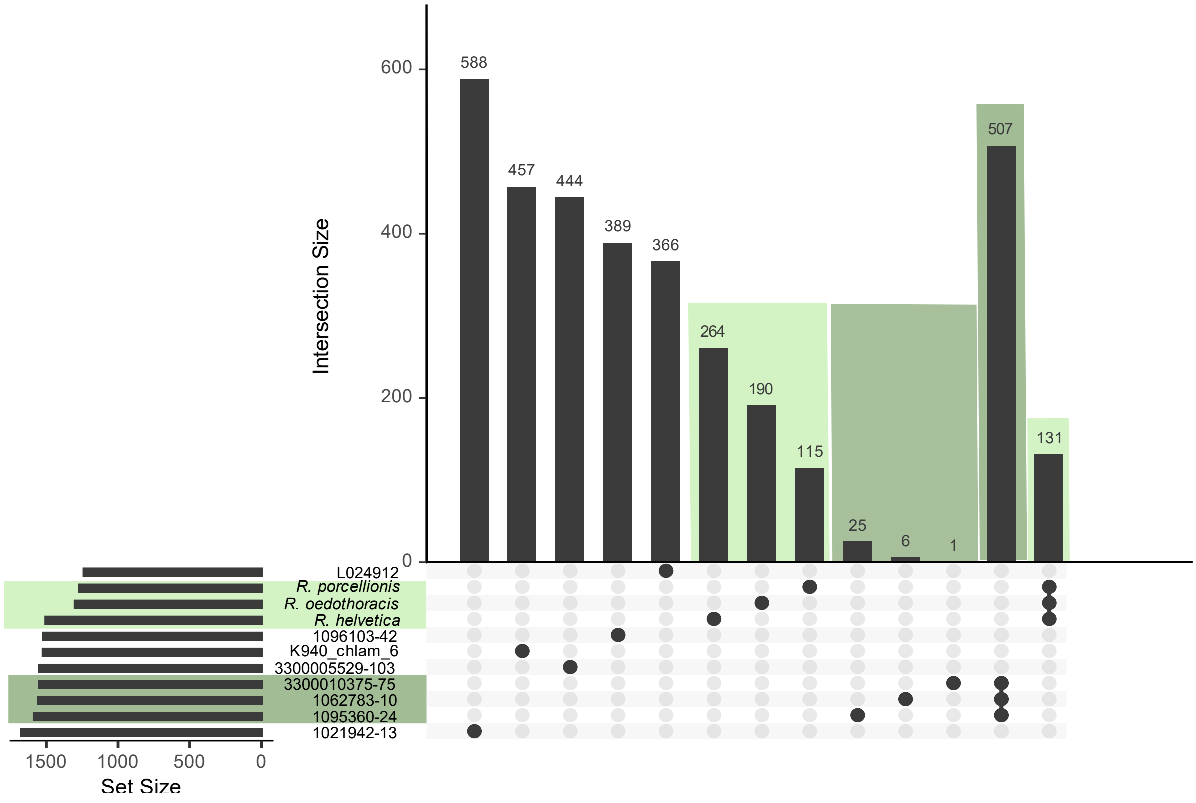


**Supplementary figure 4:** Pangenome of Rhabdochlamydiaceae. Pangenomes were visualized by UpsetR (v1.4.0) in R (v4.0.3). Genomes belonging to one genus are represented by the same color. Genomes belonging to the genus *Rhabdochlamydia* are colored in light green; another genus consisting of MAGs is labelled in dark green. The set size is the total number of OGs (including eggNOG and de novo clustered OGs) found in a certain genome. Black dots represent genomes and lines represent shared OGs. The intersection size represents the number of OGs occurring in the respective group of genomes.

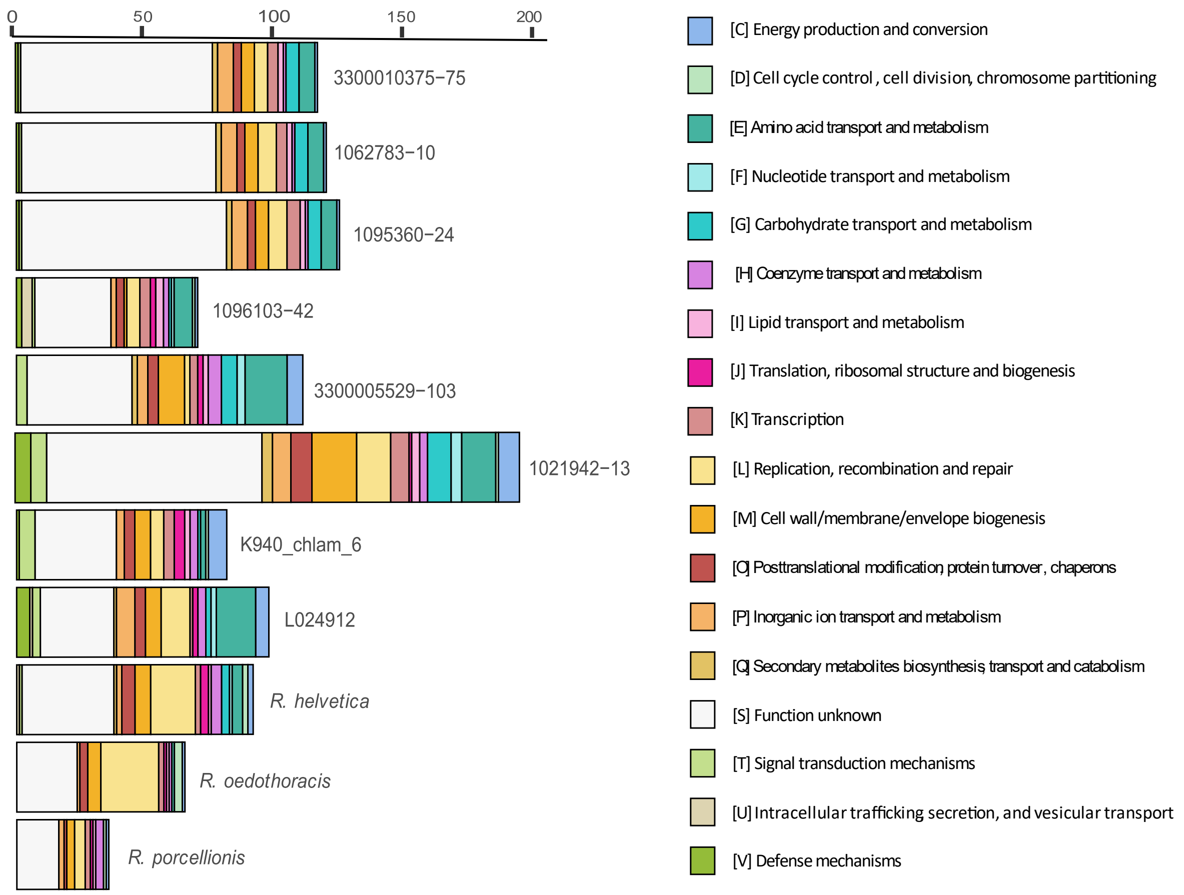


**Supplementary figure 5: Stacked-bar chart showing functional categories of rhabdochlamydiae accessory genomes.** The analysis only includes genes with a known function. For the plot only unambiguous annotations were used. For genomes belonging to the same genus i.e., the first three and the last three genomes the genus-specific genes were added to the accessory genomes.

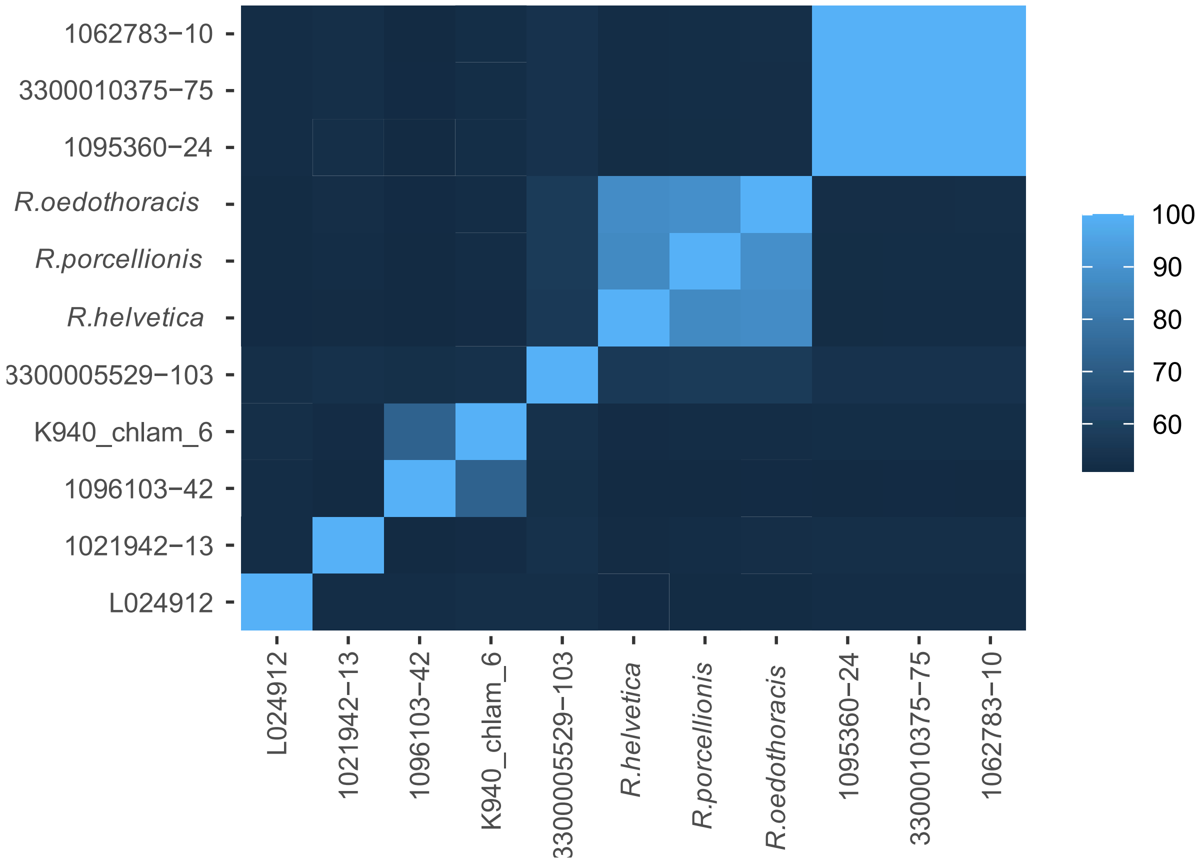


**Supplementary figure 6:** Average amino acid identity (AAI) of rhabdochlamydiae genomes. The two-way AAI was calculated for each pair of genomes belonging to the family Rhabdochlamydiaceae as described in Konstantinidis et al. (2005) [7]. For all proteins of the query genome a reciprocal best-match was searched in the reference genome. The AAI was calculated considering only those orthologous proteins. We regarded two genomes to belong to one genus if AAI >= 80 %.

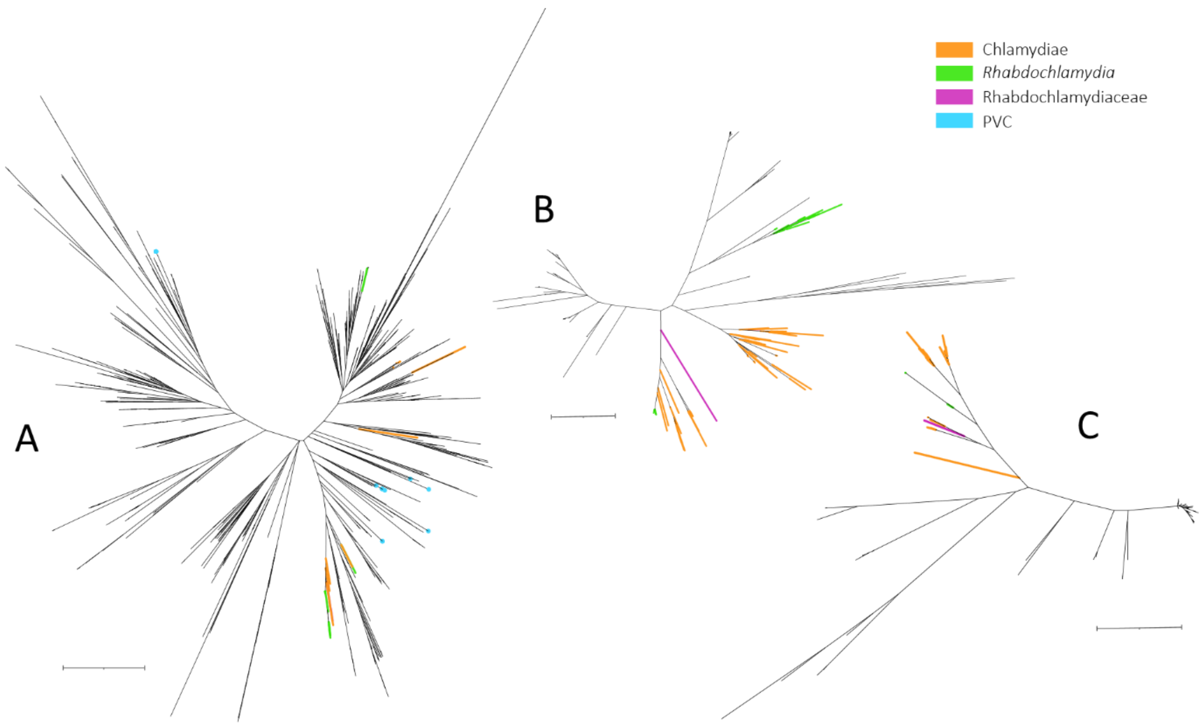


**Supplementary figure 7:** Phylogeny of the three most abundant functional transposases of *R. oedothoracis*. (**A**) Gene tree for transposase ISRhOegibbosus1 (IS30 family); (**B**) Gene tree for transposase ISRhOegibbosus2 (IS630 family); (**C**) Gene tree for transposase ISRhOegibbosus6 (IS630 family). Chlamydial transposases are indicated by orange, green, and pink branches; blue branches represent homologs from other members of the PVC phylum and black branches represent homologs from other bacteria. Gene trees were calculated as described in Koestlbacher et al. (2021) [8]. Trees were visualized and edited using the Interactive Tree Of Life [9]. As there is no information about the completeness of the transposase genes in the other genomes, the phylogenetic placement may be biased in some cases. Tree scale: 1.
