## Supplementary Tables for "Ecology and evolution of chlamydial symbionts of arthropods"

**Supplementary table S1: Basic dataset info.**

| Name | Identifier | Accession | IMG genome ID | CDS | GC [%] | Genome size [bp] | Family | Completness [%] | Heterogeneity [%] | Contamination [%] |
|---|---|---|---|---|---|---|---|---|---|---|
| Chlamydia abortus S26/3 | CHLAB | GCF_000026025.1 | | 935 | 39.87 | 1144377 | Chlamydiaceae | 98.28 | 0.00 | 0.00 |
| Chlamydia ibidis 10-1398/6 | CHLAIB | GCF_000454725.1 | | 939 | 38.32 | 1146066 | Chlamydiaceae | 94.83 | 0.00 | 0.00 |
| Chlamydia sp. RSHA | CHLBUT | GCF_900634605.1 | | 942 | 38.29 | 1146546 | Chlamydiaceae | 98.28 | 0.00 | 0.00 |
| Candidatus Chlamydia corallus strain G3/2742-324 | CHLCOR | GCF_002817655.1 | | 1016 | 39.28 | 1203974 | Chlamydiaceae | 98.28 | 0.00 | 0.00 |
| Chlamydophila caviae GPIC | CHLCV | GCF_000007605.1 | | 976 | 39.19 | 1181356 | Chlamydiaceae | 98.28 | 0.00 | 0.00 |
| Chlamydia felis Fe/C-56 | CHLFF | GCF_000009945.1 | | 974 | 39.34 | 1173791 | Chlamydiaceae | 98.28 | 0.00 | 0.00 |
| Chlamydia gallinacea 08-1274/3 | CHLGAL | GCF_000471025.2 | | 912 | 37.9 | 1067202 | Chlamydiaceae | 94.83 | 0.00 | 0.00 |
| Chlamydia muridarum str. Nigg | CHLMU | GCF_000006685.1 | | 908 | 40.31 | 1080451 | Chlamydiaceae | 98.28 | 0.00 | 0.00 |
| Chlamydia psittaci 6BC | CHLP6 | GCF_000204255.1 | | 988 | 39.02 | 1179220 | Chlamydiaceae | 98.28 | 0.00 | 0.00 |
| Chlamydophila pecorum E58 | CHLPE | GCF_000204135.1 | | 934 | 41.08 | 1106197 | Chlamydiaceae | 100.00 | 0.00 | 0.00 |
| Chlamydophila pneumoniae CWL029 | CHLPN | GCF_000008745.1 | | 1029 | 40.58 | 1230230 | Chlamydiaceae | 98.28 | 0.00 | 0.00 |
| Chlamydophila pneumoniae LPCoLN | CHLPP | GCF_000024145.1 | | 1105 | 40.5 | 1248550 | Chlamydiaceae | 98.28 | 0.00 | 0.00 |
| Chlamydia sp. 2742-308 | CHLSAN | GCF_001653975.1 | | 940 | 38.5 | 1120737 | Chlamydiaceae | 98.28 | 0.00 | 0.00 |
| Chlamydia suis MD56 | CHLSUIS | GCF_000493885.1 | | 892 | 42.01 | 1079129 | Chlamydiaceae | 98.28 | 0.00 | 0.00 |
| Chlamydia trachomatis 434/Bu | CHLT2 | GCF_000068585.1 | | 880 | 41.33 | 1038842 | Chlamydiaceae | 98.28 | 0.00 | 0.00 |
| Chlamydia trachomatis A/HAR-13 | CHLTA | GCF_000012125.1 | | 900 | 41.27 | 1051969 | Chlamydiaceae | 96.55 | 0.00 | 0.00 |
| Chlamydia trachomatis D/UW-3/CX | CHLTR | GCF_000008725.1 | | 887 | 41.31 | 1042519 | Chlamydiaceae | 96.55 | 0.00 | 0.00 |
| 1041544-29 | 1041544-29 | | 1041544 | 2382 | 38.09 | 2850003 | Parachlamydiaceae | 98.28 | 0.00 | 0.00 |
| 3300009084-136 | 3300009084-136 | | 3300009084 | 1941 | 34.86 | 2250271 | Parachlamydiaceae | 96.55 | 0.00 | 0.00 |
| 3300009084-150 | 3300009084-150 | | 3300009084 | 1747 | 45.72 | 1750468 | Parachlamydiaceae | 100 | 0.00 | 0.00 |
| Protochlamydia sp. ACF82 | ACE82 | JAEMUB000000000 | | 2233 | 34.49 | 2331284 | Parachlamydiaceae | 95.69 | 0.00 | 0.00 |
| Parachlamydia sp. ACF125 | ACF125 | JAEMUD000000000 | | 1919 | 40.77 | 2584652 | Parachlamydiaceae | 98.28 | 0.00 | 0.00 |
| Chlamydia sp. 32-24 | CHL3224 | GCA_001897185.1 | | 2075 | 32.42 | 2529957 | Parachlamydiaceae | 98.28 | 0.00 | 0.00 |
| Chlamydiales bacterium 38-26 | CHL3826 | GCA_001897225.1 | | 2327 | 38.12 | 2834110 | Parachlamydiaceae | 98.28 | 0.00 | 0.00 |
| Protochlamyida sp. EI2 | EI2 | GCA_000813625.1 | | 2150 | 34.82 | 2397675 | Parachlamydiaceae | 96.55 | 0.00 | 0.00 |
| Neochlamydia sp. EPS4 | EPS4 | GCF_000813665.1 | | 1882 | 38.09 | 2530677 | Parachlamydiaceae | 96.55 | 0.00 | 0.00 |
| Chlamydiales bacterium STE3 | HSC3 | VKHO00000000 | | 2176 | 39.17 | 2231767 | Parachlamydiaceae | 98.28 | 0.00 | 0.00 |
| Parachlamydia acanthamoebae OEW1 | OEW1 | GCA_000812225.1 | | 2309 | 39.04 | 3008885 | Parachlamydiaceae | 94.83 | 0.00 | 0.00 |
| Parachlamydia acanthamoebae UV-7 | PARAV | GCF_000253035.1 | | 2532 | 39.04 | 3072383 | Parachlamydiaceae | 96.55 | 0.00 | 0.00 |
| Candidatus Protochlamydia amoebophila UWE25 | PARUW | GCF_000011565.1 | | 1841 | 34.72 | 2414465 | Parachlamydiaceae | 98.28 | 0.00 | 0.00 |
| Parachlamydia sp. C2 | PROGREC2 | GCA_001545115.1 | | 2766 | 42.05 | 3424182 | Parachlamydiaceae | 100.00 | 0.00 | 0.00 |
| Protochlamydia naegleriophila | ProNeg | GCA_001499655.1 | | 2520 | 42.44 | 3030375 | Parachlamydiaceae | 100.00 | 0.00 | 0.00 |
| Protochlamydia massiliensis | PRONEGDIA | GCF_000751535.1 | | 2451 | 42.75 | 2956128 | Parachlamydiaceae | 98.28 | 0.00 | 0.00 |
| Parachlamydia sp. isolate BC.030 | PROPHO | GCA_002786175.1 | | 2540 | 41.53 | 3042961 | Parachlamydiaceae | 100.00 | 0.00 | 0.00 |
| Rubidus massiliensis | Rubis | GCA_000756735.1 | | 2446 | 32.64 | 2821221 | Parachlamydiaceae | 98.28 | 0.00 | 0.00 |
| Neochlamydia sp. TUME1 | TUME1 | GCF_000813645.1 | | 1879 | 38.02 | 2546323 | Parachlamydiaceae | 96.08 | 0.00 | 0.00 |
| 1021942-13 | 1021942-13 | | 1021942 | 1792 | 42.3 | 2050771 | Rhabdochlamydiaceae | 97.41 | 0.00 | 0.00 |
| 1062783-10 | 1062783-10 | | 1062783 | 1724 | 45.19 | 1859781 | Rhabdochlamydiaceae | 100 | 0.00 | 0.00 |
| 1095360-24 | 1095360-24 | | 1095360 | 1750 | 45.12 | 1889532 | Rhabdochlamydiaceae | 100 | 0.00 | 0.00 |
| 1096103-42 | 1096103-42 | | 1096103 | 1637 | 44.67 | 1644418 | Rhabdochlamydiaceae | 95.69 | 0.00 | 0.00 |
| 3300005529-103 | 3300005529-103 | | 3300005529 | 1695 | 43.86 | 1937521 | Rhabdochlamydiaceae | 100 | 0.00 | 0.00 |
| 3300010375-75 | 3300010375-75 | | 3300010375 | 1704 | 45.18 | 1846245 | Rhabdochlamydiaceae | 94.59 | 0.00 | 0.00 |
| Chlamydiae bacterium K940_chlam_6 | K940_chlam_6 | GCA_001796315.1 | | 1750 | 42.93 | 1505150 | Rhabdochlamydiaceae | 95.69 | 0.00 | 0.00 |
| Chlamydiae bacterium RIFCSPLOWO2_02_FULL_49_12 | L024912 | from PMID: 32142706 | | 1313 | 48.99 | 1413529 | Rhabdochlamydiaceae | 96.55 | 0.00 | 0.00 |
| Rhabdochlamydia helvetica | RhabHel | from PMID: 30949677 | | 1717 | 36.2 | 1854477 | Rhabdochlamydiaceae | 94.59 | 0.00 | 0.00 |

**Supplementary table S2: Basic genome statistics of complete *Rhabdochlamydia* genomes.**

| | ***R. porcellionis* 15C** | ***R. oedothoracis* W744** |
|---|---|---|
| Closed | yes | yes |
| Estimated size [mb] | 1.49 | 1.88 |
| Plasmid [kb] | 19.7 | 38.9 |
| Contigs | 2 | 2 |
| Completeness [%] | 100 | 100 |
| Contamination [%] | 0 | 0 |
| Heterogeneity [%] | 0 | 0 |
| tRNA | 37 | 37 |
| rRNA | 6 | 6 |
| CDS | 1.348 | 1.569 |
| GC [%] | 35.4 | 36.2 |

**Supplementary table S3: OGs discriminating the genus *Rhabdochlamydia* from other members of the family Rhabdochlamydiaceae**

| OG missing in genomes of the family Rhabdochlamydiaceae but present in all members of the genus *Rhabdochlamydia* | Functional Category | Membrane [Y/N] | Description | Evidence |
|---|---|---|---|---|
| 05CPH | H | N | Catalyzes the formation of S-adenosylmethionine from methionine and ATP | |
| 05DV8 | S | N | filamentation induced by cAMP protein Fic | |
| 05EGX | M | Y | Nad-dependent epimerase dehydratase | Functional Category |
| 05ERD | L | N | Involved in base excision repair of DNA damaged by oxidation or by mutagenic agents. Acts as DNA glycosylase that recognizes and removes damaged bases. Has a preference for oxidized purines, such as 7,8-dihydro-8-oxoguanine (8-oxoG). Has AP (apurinic apyrimidinic) lyase activity and introduces nicks in the DNA strand. Cleaves the DNA backbone by beta-delta elimination to generate a single-strand break at the site of the removed base with both 3'- and 5'-phosphates (By similarity) | |
| 05F0X | D | N | Cobyrinic acid ac-diamide synthase | |
| 05F23 | K | Y | Fibronectin-binding protein | https://doi.org/10.1007/s00430-019-00644-3 |
| 05RKK | S | N | Domain of unknown function (DUF202) | |
| 05Z3Q | J | N | tRNA rRNA methyltransferase | |
| 06A7V | S | Y | Permease of the drug metabolite transporter DMT superfamily | GO Term: Cellular Component |
| 06VZX | L | N | establishment of viral latency | |
| 071KN | S | Y | MOMP-like family | https://doi.org/10.1128/IAI.69.5.3082-3091.2001 |
| 07EW0 | M | Y | peptidase | Functional Category |
| 07GNU | C | Y | TLC ATP/ADP transporter | GO Term: Cellular Component |
| 07SJN | I | N | Catalyzes the phosphorylation of the position 2 hydroxy group of 4-diphosphocytidyl-2C-methyl-D-erythritol (By similarity) | |
| 07WZ0 | P | N | ferritin dps family protein | |
| 085BK | S | N | UPF0109 protein | |
| 08JHT | E, G | Y | EamA-like transporter family | GO Term: Cellular Component |
| 08MMU | S | N | Pfam:DUF2843 | |
| 08NMA | L | N | Plasmid and phage replicative helicase | |
| 08SEI | S | N | SOUL heme-binding protein | |
| 0NHB5 | O | N | PPIases accelerate the folding of proteins | |
| 0QNFT | M | Y | glycosyl transferase group 1 | Functional Category |
| 05ZQP | S | N | NA | |
| 060UR | S | N | NA | |

| OG present in all genomes of the family Rhabdochlamydiaceae but completely missing in the genus *Rhabdochlamydia* | Functional Category | Membrane [Y/N] | Description | Evidence |
|---|---|---|---|---|
| 05C5C | M | Y | n-acetylmuramoyl-l-alanine amidase | Functional Category |
| 05C7Z | O | N | Conserved Protein | |
| 05DCI | S | N | Phosphohydrolase | |
| 05DKP | M | Y | glycosyl transferase, family 9 | Functional Category |
| 05VH8 | J | N | Binds as a heterodimer with protein S6 to the central domain of the 16S rRNA, where it helps stabilize the platform of the 30S subunit (By similarity) | |
| 07NVQ | N | Y | Flagellar biosynthesis protein, FliO | Functional Category |
| 07RI4 | M | Y | Transfers the fatty acyl group on membrane lipoproteins (By similarity) | Functional Category |
| 08RMU | P | N | ferritin dps family protein | |

**Supplementary table S4: Overview of transposases of *R. porcellionis* and *R. oedothoracis*. For each transposon the counts of functional genes and pseudogenes are depicted. The length always refers to functional genes.**

| Organism | Name, IS group/family | # pseudogenes | # functional genes | # total | Length [nt] | Plasmid [Y/N] |
|---|---|---|---|---|---|---|
| *R. oedothoracis* | Transposase ISRhOegibbosus1, IS30 family | 25 | 74 | 99 | 1.845 | Y |
| *R. oedothoracis* | Transposase ISRhOegibbosus2, IS630 family | 42 | 42 | 84 | 1.035 | Y |
| *R. oedothoracis* | Transposase ISRhOegibbosus3, IS1031 group IS5 family | 79 | 0 | 79 | | N |
| *R. oedothoracis* | Transposase ISRhOegibbosus4, IS1634 family | 3 | 0 | 3 | | N |
| *R. oedothoracis* | Transposase ISRhOegibbosus5, IS427 group IS5 family | 32 | 0 | 32 | | Y |
| *R. oedothoracis* | Transposase ISRhOegibbosus6, IS630 family | 31 | 9 | 40 | 804 | Y |
| *R. oedothoracis* | Transposase ISRhOegibbosus7, IS110 group IS1111 | 18 | 0 | 18 | | N |
| *R. oedothoracis* | Transposase ISRhOegibbosus8, ISL2 group IS5 family | 17 | 0 | 17 | | N |
| *R. oedothoracis* | Transposase ISRhOegibbosus9, IS630 family | 11 | 2 | 13 | 1.032 | N |
| *R. oedothoracis* | Transposase ISRhOegibbosus10, IS481 family | 12 | 0 | 12 | | Y |
| *R. oedothoracis* | Transposase ISRhOegibbosus11, IS982 family | 3 | 0 | 3 | | N |
| *R. oedothoracis* | Transposase ISRhOegibbosus12, ISAs1 family | 2 | 0 | 2 | | N |
| *R. oedothoracis* | Transposase ISRhOegibbosus13, IS630 family | 3 | 0 | 3 | | N |
| *R. oedothoracis* | Transposase ISRhOegibbosus14, ISAs1 family | 7 | 1 | 8 | 1.143 | N |
| *R. oedothoracis* | Transposase ISRhOegibbosus15, IS3 group IS3 family | 1 | 0 | 1 | | N |
| *R. oedothoracis* | Transposase ISRhOegibbosus16, IS481 family | 1 | 0 | 1 | | N |
| *R. procellionis* | Transposase ISRhPorc1, ISL2 group IS5 family | 2 | 0 | 2 | | N |
| *R. procellionis* | Transposase ISRhPorc2, IS630 family | 12 | 0 | 12 | | N |
| *R. procellionis* | Transposase ISRhPorc3, ISL2 group IS5 family | 5 | 0 | 5 | | N |
| *R. procellionis* | Transposase, PD-(D/E)XK nuclease family | 0 | 1 | 1 | 936 | N |
